# Kinomorphs: Shape-shifting tissues for developmental engineering

**DOI:** 10.1101/768218

**Authors:** John M. Viola, Catherine M. Porter, Ananya Gupta, Mariia Alibekova, Louis S. Prahl, Alex J. Hughes

## Abstract

Current methods for building tissues usually start with a non-biological blueprint, or rely on self-organization, which does not extend to organ-scales. This has limited the construction of large tissues that simultaneously encode fine-scale cell organization. Here we bridge scales by mimicking developmental dynamics using “kinomorphs”, tissue scaffolds that undergo globally programmed shape and density changes to trigger local self-organization of cells in many locations at once. In this first report, we focus on mimicking the extracellular matrix (ECM) compaction and division into leaflets that occurs in kidney collecting duct development. We start by creating single-cell resolution cell patterns in ECM-mimetic hydrogels that are >10x larger than previously described, by leveraging photo-lithographic technology. These patterns are designed to mimic the branch geometry of the embryonic kidney collecting duct tree. We then predict the shape dynamics of kinomorphs driven by cell contractility-based compaction of the ECM using kinematic origami simulations. We show that these dynamics spur centimeter-scale assembly of structurally mature ~50 μm-diameter epithelial tubules that are locally self-organized, but globally programmed. Our approach prescribes tubule network geometry at ~5x smaller length-scales than currently possible using 3D printing, and at local cell densities comparable to *in vivo* tissues. Kinomorphs could be used to scaffold and “plumb” arrays of organoids in the future, by guiding the morphogenesis of epithelial networks. Such hybrid globally programmed/locally self-organized tissues address a major gap in our ability to recapitulate organ-scale tissue structure.

**Significance Statement:** Engineers are attempting to build tissues that mimic human diseases outside of the body. Although stem cells can be coaxed to form small organoids with a diversity of cell types, they do not properly organize over large distances by themselves. We report a strategy to mimic developmental processes using dynamic materials that attempt to guide a cellular “blueprint” towards a more complex tissue endpoint. We call these materials kinomorphs, combining the Greek kinó (propel, drive) and morfí (form, shape), since they seek to shepherd both the shape and developmental trajectory of cell collectives within them. Kinomorphs could pave the way towards organ-scale synthetic tissues built through a hybrid of engineering and self-organization strategies.

## Introduction

Tissue morphogenesis builds structural features of organisms through changes in cell and ECM position, density, and composition over time. Engineers are attempting to control these processes to build more life-like tissues using microfluidic, 3D printing, and organoid technologies (1–4). Organoids, 3D tissues grown from stem cells, have become an essential route because of the remarkable cellular diversity and spatial structure that can be achieved through spontaneous cell sorting and spatially restricted differentiation (5). However, organoids have largely not addressed longer length-scale (>0.5 mm) tissue organization processes beyond local self-organization. More prescriptive tissue engineering scaffolds built through 3D printing of cells and ECM, or subtractive hollowing of lumens in hydrogel using “fugitive inks” can impose such long-range cues (6–9). However, tissues made using these approaches are often not cell dense, do not reconstruct critical tissue interfaces like the epithelial-mesenchymal interface, are not intended to mimic tissue dynamics, and/or do not achieve ECM densities of *in vivo* tissues.

For example in the kidney, organoid models can currently produce nephron-like multicellular collectives (10), but not their full connectivity via a contiguous, tree-like collecting duct network that spans centimeters. Recombination of mouse embryonic stem cell-derived ureteric, nephron, and fetal stromal progenitor cells was found to recapitulate collecting duct branching to 3-4 branch generations in a ~1 mm-diameter organoid, and some branching activity was also seen in an analogous experiment using human induced pluripotent stem cells (11). However, extending these exciting results to length-scales approaching those of whole organs will likely require new engineering approaches to control developmental patterning cues, starting in a well-studied prototype model.

One such model is the kidney tubule-like cell line family known as Madin-Darby canine kidney cells (MDCKs). MDCKs are an attractive starting point because they can form single-cell layer spheroids and randomly oriented tubules that lumenize spontaneously in different 3D culture conditions (12, 13). Despite the selforganization properties of MDCKs, there is little conceptual framework for orchestrating structure over longer distances and set geometries. We propose that dynamic tissue scaffolds that mimic morphogenetic motifs may be a route towards achieving more structurally complex endpoints.

Indeed, there are several such motifs in collecting duct morphogenesis, in which the ureteric bud epithelium invades the surrounding mesenchyme (an initially loose cell/ECM composite). Aspects of these are also shared with other organ systems including the lung, mammary gland, and salivary gland (14). These motifs include duct elongation, branching, thinning/narrowing, and mutual avoidance, which prevents fusion of neighboring ducts (15, 16). The overall effect of this is to compact and divide the mesenchyme ECM into leaflets. These “leafleting” and compaction dynamics have a distinct geometric analogy with folding processes that are potentially open to engineering control (Movie S1).

To autonomously achieve sufficient control over ECM leafleting and compaction dynamics within a tissue scaffold requires tight “bottom-up” control. One way to achieve this is to precisely determine the starting composition and geometry of scaffolds containing cells and ECM. In previous work we developed a DNA-patterned assembly of cells (DPAC) approach. Cell populations are labeled with lipid-modified ssDNA oligos that passively insert into cell membranes (17), and then temporarily adhered by base pairing to spots of a complementary ssDNA strand patterned onto a glass slide (18, 19). DPAC has several advantages over other cell patterning methods (20, 21), namely that different cell populations can be independently patterned in the same experiment using orthogonal pairs of oligo sequences, cells can be patterned with spatial resolution on the order of 10 μm, and cell patterns can be transferred from the assembly interface into any of a range of hydrogels solidified around them. However, DPAC is significantly limited by the speed at which oligo features can be printed, and thus the scale at which tissue scaffolds can be built.

Given an ability to set a particular blueprint using bottom-up cell and ECM patterning, we also require a “top-down” framework that would predict the relationship between single-cell dynamics and the dynamics of the scaffold as a whole. Perhaps the best understood framework that links 2D blueprints to 3D dynamics is the field of origami, where out-of-plane 3D shape change is determined by material strains at crease networks (22). In previous work we found that mechanical strains caused by interactions between cells (especially contractile cell types) and ECM on the top interface of hydrogel sheets were relieved by the formation of negative curvatures (valleys), whereas strains on the bottom interface led to positive curvatures (mountains) (19). Further, we found that human umbilical vein endothelial cells directionally migrated along folds in ECM sheets in response to these out-of-plane curvature programs generated by contractile fibroblasts (19). These dynamics point towards the possibility of imposing a top-down shape change that would transform a bottom-up cell blueprint into a programmed tissue output.

In this paper, we seek to 1) vastly increase the scale of cell patterning in ECM sheets, so that we can 2) mimic ECM dynamics consistent with kidney development to guide the formation of kidney tubules across cm scales. Using high-throughput, high-resolution cell patterning, we build kinomorphs – cell-ECM composite sheets that undergo a prescribed trajectory in shape and size/density. This trajectory creates niches for kidney tubule tracts that promote structural maturation, as defined by polarity and lumenization. Kinomorphs are a promising approach for achieving mechano-geometric control over cell collectives, with future applications in meso-scale scaffolding of organoids and other efforts to mimic dynamic developmental processes.

## Results

In kidney collecting duct network morphogenesis, tubules branch and extend, thereby compacting the surrounding mesenchyme with an estimated area strain of −20% over 70 hr in explant cultures of E11.5 mouse kidneys (Fig. 1A). We first asked whether mimicking these ECM compaction and leafleting dynamics were sufficient to guide the *in vitro* morphogenesis of MDCK tubules. Culturing groups of MDCKs assembled just below the interface of collagen I-Matrigel hydrogels using DPAC caused them to condense into clusters with smooth boundaries (Fig. 1B). Subsequent immunostaining of these clusters revealed proper localization of F-actin to apical membranes, and E-cadherin to lateral cell-cell contacts – hallmarks of epithelial polarization (23). These spheroids formed when the hydrogel was adhered to an underlying substrate. However, if the hydrogel was allowed to float in medium, spheroids tended to elongate and fuse to form tubule-like structures near the curled and compacted edges of the gel. Live imaging showed that these dynamics occurred through collective cell movements, remodeling of surrounding ECM fibers by cell tractions, and fusion between neighboring cell clusters (Movie S2). Based on this, we concluded that tubule formation appeared to be permitted in our system by gel compaction at curled creases. We hypothesized that bringing this property under geometric control could allow us to build tubule networks with defined geometries and connectivities. In short, since creases guided MDCK tubule formation, we reasoned that building crease networks would guide tubule network formation.

**Fig. 1:**
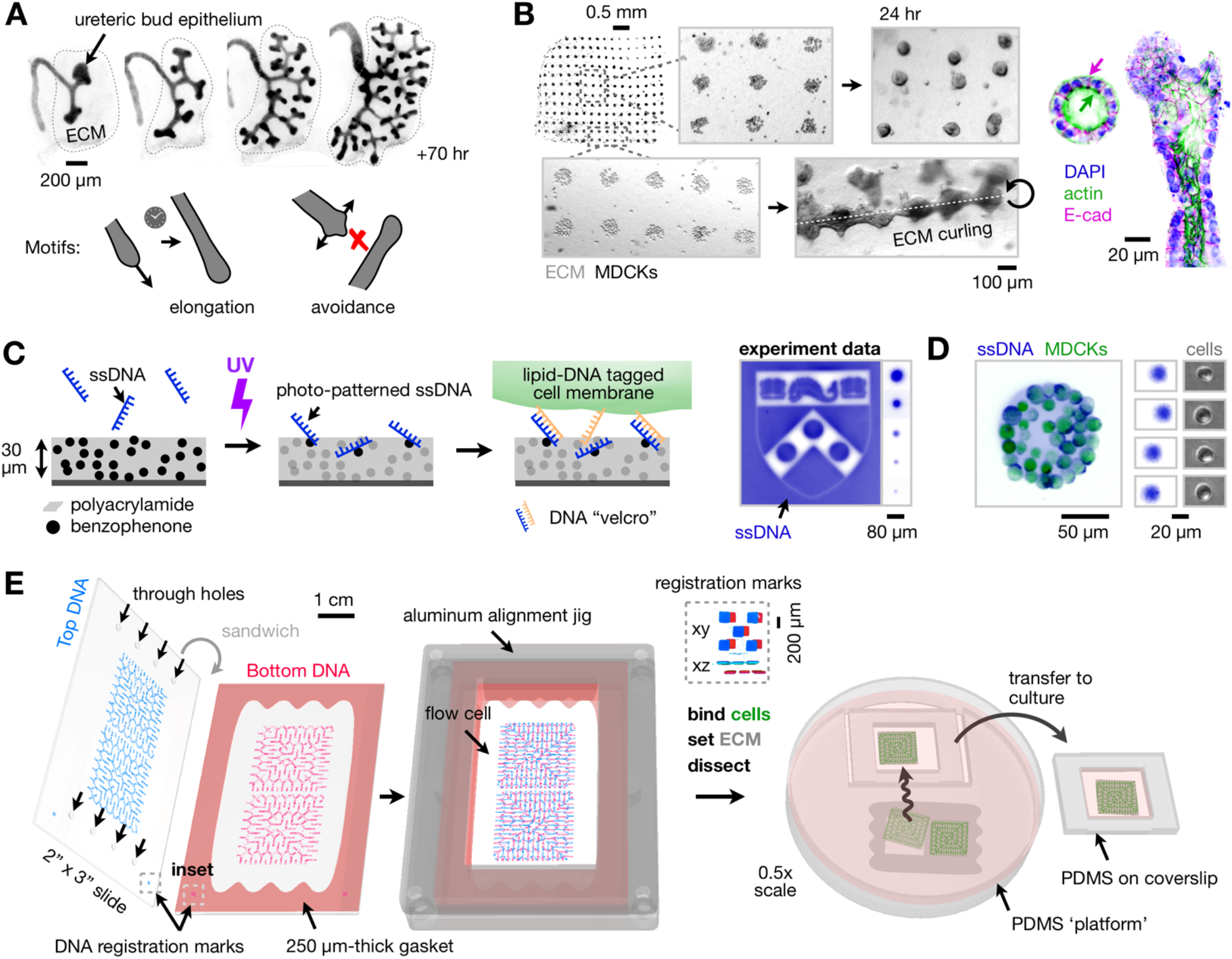
Mimicking kidney collecting duct morphogenesis through fusion of photopatterned cell clusters in pliable ECM gels. (**A**) Time-lapse fluorescence micrographs of embryonic day 11.5 kidney ureteric bud branching into the metanephric mesenchyme (modified from (30)). Tubule branching, elongation, and avoidance processes separate and compact the mesenchyme into leaflets. (**B**) *Left*, phase micrographs of an array of Madin-Darby canine kidney (MDCK) clusters patterned using photo-lithographic DNA-patterned assembly of cells (pDPAC) and embedded just under the surface of a 250 μm-thick collagen I-Matrigel sheet. Clusters form spheroids that fuse at the pliable edges of the sheet. *Right*, Immunofluorescence micrographs of spheroids and fused tubule structures, showing apical F-actin distribution (green arrow), and lateral E-cadherin distribution (magenta arrow). (**C**) *Left*, pDPAC workflow showing ssDNA patterning onto a thin benzophenone-functionalized gel attached to glass. *Right*, fluorescence micrograph of a gel-bound SYBR Gold-labeled DNA pattern and spots of different diameters, down to 10 μm. (**D**) *Left*, Fluorescence micrograph of SYBR Gold-labeled live MDCKs and an underlying F DNA spot. Cells were pre-labeled with complementary F’ lipid-DNA and CellTracker dye. Note plasma membrane localization of SYBR signal associated with lipid-DNA. *Right*, SYBR Gold-labeled 20 μm diameter DNA spots and single MDCK cells attached to such spots. (**E**) Schematic of microfluidic flow cell assembly using pDPAC slides separated by a gasket, cell capture, hydrogel embedding, and transfer into culture. *Inset*, SYBR Gold-labeled registration marks used for aligning DNA patterns (*SI Appendix*).

With this potential control strategy emerging, we first sought to make significant improvements in the length-scale of our cell patterning capabilities in order to match those relevant to kidney branching morphogenesis. The previous microcantilever-based printing method used to deposit ssDNAs in DPAC is limited in throughput and spatial scale, since individual DNA spots can be printed with a frequency of only ~1 Hz. We reasoned that we could lift this limitation by shifting to a photo-lithography approach (“pDPAC”) that would enable millions of DNA features to be printed simultaneously rather than serially.

We approached this by repurposing a photo-active biomolecule patterning technology that we previously developed for the immobilization of proteins within polyacrylamide gels (24). We first polymerized a 30 μm-thick sheet of 4% polyacrylamide gel containing a photo-reactive benzophenone-methacrylamide co-monomer (Fig. 1C). Applying 254 nm light through photomasks then allowed us to tether unmodified ssDNA oligos onto and within the gel in feature sizes down to 10 μm (actual: 11.1 μm ± 5.8% CV, *n* = 5). These features were sufficient to direct the adhesion of single cells labeled with a complementary lipid-DNA to the polyacrylamide surface (Fig. 1D) with relatively low binding of cells to unpatterned areas (~4 cells mm^-2^). We use single-letter nicknames to denote different patterned DNA/lipid-DNA strand pairs here – in this case “F” was patterned onto the pDPAC substrate and cells were labeled with F’, the reverse complement of F (see *SI Appendix* for full sequences). We found that both the amount of DNA patterned and the subsequent cell capture efficiency onto pDPAC slides increased as a function of the UV dose (Fig. S1A). Further, thymine bases were primarily responsible for DNA patterning onto pDPAC gels (Fig. S1B). We therefore included a polyT20 tail on DNAs to sensitize them to immobilization during UV exposure. Patterning each DNA strand could be performed in roughly 45 min at scales of up to 5.1 cm x 7.6 cm (or 3.9 x 10^7^ features capable of patterning a single cell), resulting in a speed-up of >50x over the previous printing method. The hydrophilic polyacrylamide patterning interface also directly enabled large-scale embedding of cell patterns into collagen I-Matrigel hydrogels, since these gels showed very little adhesion to polyacrylamide during transfer into culture.

With a large-scale bottom-up fabrication method in place, we reasoned that we could apply it to encode networks of creases that would each guide an MDCK tubule along it. To do this, we patterned hydrogel sheets with contractile 3T3 mouse embryonic fibroblast cells that would roughly mimic the branch geometry of the embryonic ureteric tubule network. The use of fibroblasts would enable us to achieve tight geometric control over ECM compaction by using their contractility to control where creases form, rather than relying on passive crease formation at the edges of floating gels. This would then allow us to enforce a known blueprint to the tubule self-organization process that mimics ECM leafleting and compaction dynamics in kidney epithelial branching morphogenesis. In short, we needed 1) an apparatus to encode crease networks into ECM using pDPAC, 2) a reference geometry describing the embryonic collecting duct network that would serve as a design goal, and 3) a model to predict how candidate crease networks that mimic the reference geometry would emerge from flat sheets.

For the apparatus - we created a large-format flow cell device that sandwiches two pDPAC slides at a distance of ~250 μm apart (Fig. 1E, Movie S3). Our flow cell design accommodates cell patterning on both the top and bottom polyacrylamide surfaces, from which cells are transferred into collagen 1-Matrigel after setting the gel precursor mixture within the flow cell. The hydrogel sheet is then dissected and coaxed into a culture vessel for live imaging.

Secondly, for the reference geometry - we used recently published data to simulate an example collecting duct branching pattern in embryonic day 19 (E19) mouse kidney (Fig. 2A) (25).

**Fig. 2:**
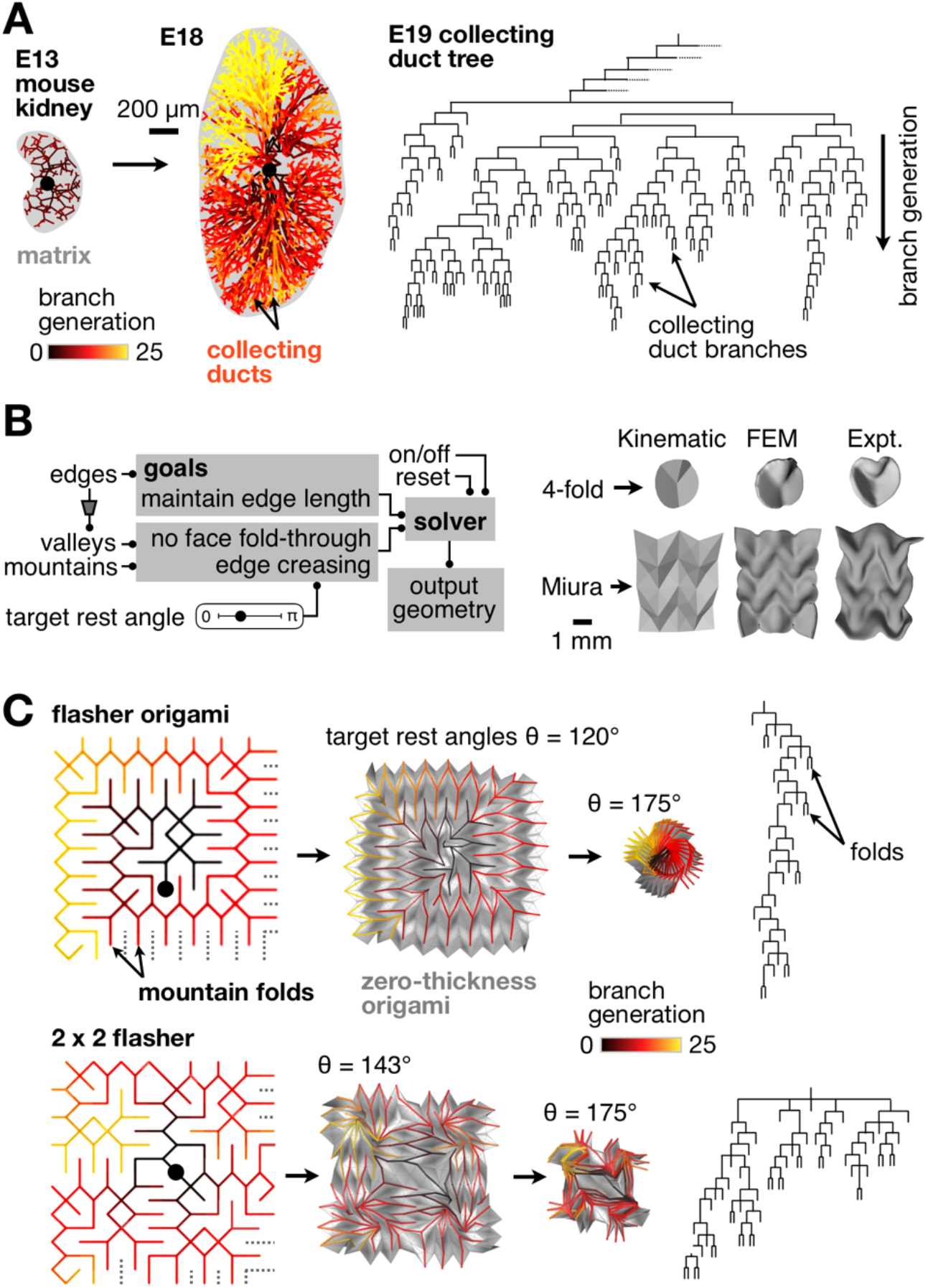
Forward-prediction of tissue structure using flasher origami crease networks that approximate kidney collecting duct geometry. (**A**) *Left*, 3D reconstructions of collecting duct geometry in embryonic mouse kidneys (“E##” = embryonic day ##), with ducts color-coded by their distance in number of branch generations from the root of the network (filled black circle, modified from (25)). *Right*, an example branch pattern simulated from experimentally measured branch probabilities (25). (**B**) *Left*, Flow chart outlining the origami simulator used to study candidate crease networks. *Right*, 3D surfaces generated from this simplified kinematic origami simulator vs. a full 3D FEM model used in previous work vs. experimentally generated self-folding ECM sheets based on classic 4-fold and Miura origami crease networks (partly reproduced from (19)). (**C**) Origami simulator output for flasher (*top*) and 2×2 tessellated flasher (*bottom*) crease networks. *Left*, crease networks for the set of mountain folds, which emanate from a single point or edge (filled black circles), color coded by the number of branch generations from a given crease to these center points. *Middle*, 3D simulator output at different target crease rest angles (θ), including the associated sheet surfaces in gray. *Right*, branch patterns describing the crease networks, ignoring the occasional short circuit.

Finally, for the model - we created a custom origami simulator to enable rapid prototyping in an environment that permits real-time investigation of the spatial transformation of candidate crease networks (Fig. 2B). We validated the simulator by measuring the “Hausdorff distance” (26) between 3D objects created with it to those generated from our previously reported FEM technique, and to those from ECM sheets folded in vitro (19) (Fig. S2). At < 50 μm on average, these distances were all well below the sheet thickness, suggesting that the much less computationally expensive origami simulator models were adequate for predicting the ~mm-to-cm scale shape dynamics of self-folding sheets.

With these three core engineering needs in place, we sought crease blueprints that approximately matched the reference branching geometry, and that would also permit approximately flat-folded creases to form throughout the design. There are some intrinsic limitations to this type of search, because arbitrary crease patterns are not guaranteed to rigidly fold into a target 3D shape or volume (22, 27). Instead, we searched existing origami designs for tree-like crease patterns that could generate networks with similar branching geometry to the embryonic kidney epithelium and modeled their folding dynamics. These efforts led us to a class of origami called “flashers” that have mountain and valley networks that each emanate from a single point or edge. Flasher networks also bifurcate with approximately similar probabilities to those observed over wide ranges in the branching hierarchy of the embryonic mouse kidney (Fig. 2C). Tessellated flasher origami designs can be generated that retain these properties up to at least 3×3 without significant modification (Fig. S3).

With a candidate crease family selected, we set about translating flasher crease networks into cell patterns. We started with patterns of 3T3 fibroblasts would actuate curvature and create crease niches to guide MDCK tubule formation in later designs. We first created “crease blocks” – sets of 20 μm-diameter ssDNA features in anisotropic grid patterns that we previously found to encode creases along specific axes within self-folding tissues (19). Each feature represents a mask position at which DNA is deposited on pDPAC slides, and thus at which cells are patterned (Fig. 3A). We then assembled a mosaic of crease blocks into the complete origami design, which we built two at a time for a total of 139,396 DNA features per experiment (equivalent to >38 hours of printing time using the original DPAC method, Movie S4). We used pDPAC to create 22.4 x 22.4 x 0.25 mm flasher kinomorphs from this design as ECM sheets with 3T3s patterned in the prospective mountain and valley networks on each side (measured thickness was 229 μm ± 12% CV for *n* = 7 sites spread across the sheet area). At 5 cm^2^, these kinomorphs were 10.5x larger in area than any self-folding tissue made previously, enabled primarily by the throughput and low adhesion advantages of pDPAC.

**Fig. 3.**
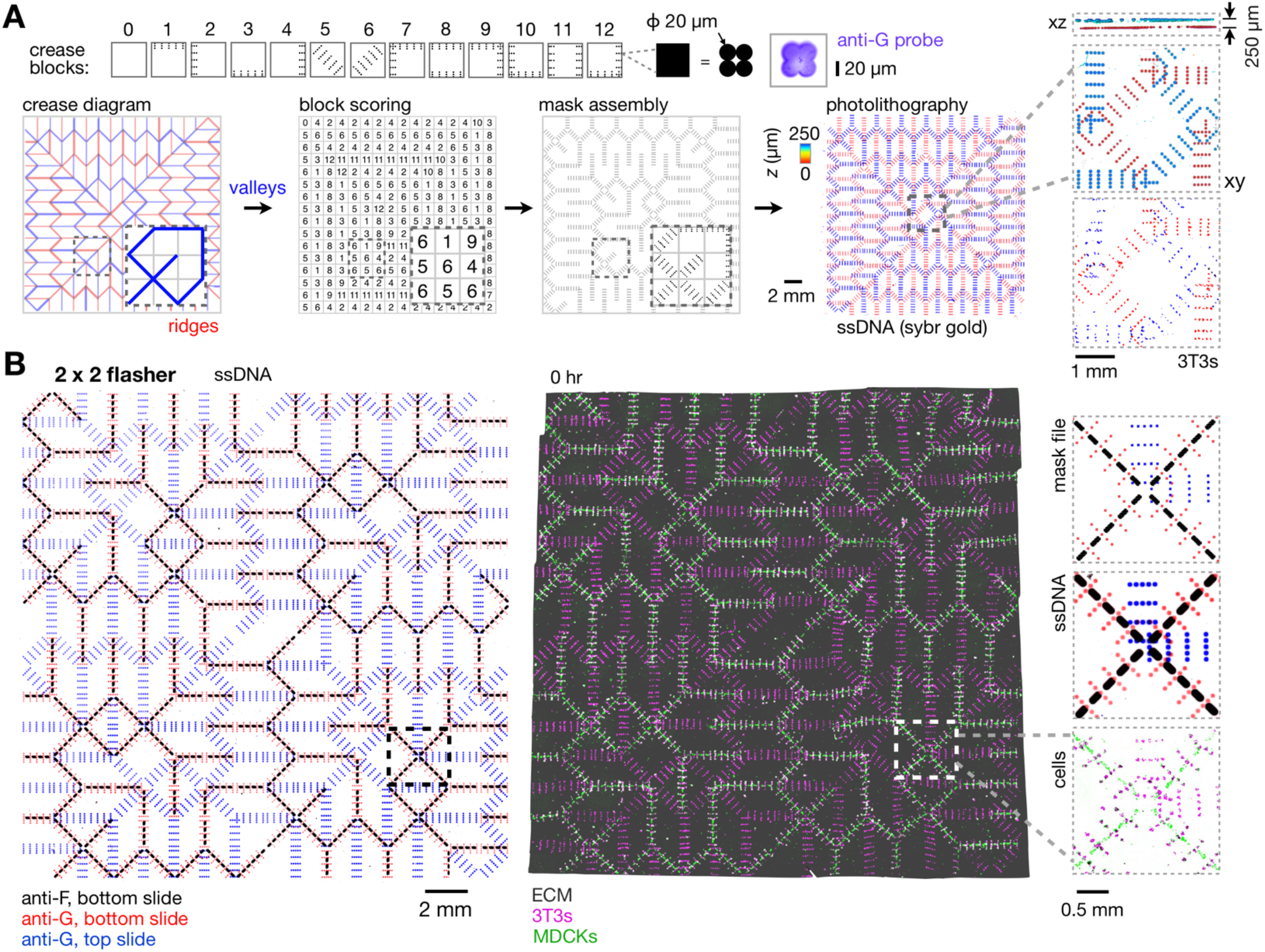
Design and construction of kinomorphs to template tissue morphogenesis at flasher creases. (**A**) *Top row*, crease blocks define pixels at which ssDNA features and thereby contractile 3T3 fibroblast cells will be placed. Each pixel encodes a 2×2 grid of 20 μm-diameter circular features. *Right*, confocal fluorescence micrograph showing G DNA bound to a pDPAC slide and stained with an anti-G-FITC probe sequence. *Bottom row*, a flasher origami crease diagram shown as valley (blue) and mountain (red) creases. Each set of creases is manually scored to associate a particular crease block. Crease blocks are then assembled into a full pixel matrix that is translated into a mask design (not shown). *At right*, confocal fluorescence micrographs of SYBR Gold-labeled DNA features on a pair of assembled pDPAC slides, color coded by depth. *Insets*, detail and xz projection of DNA features and CellTracker dye-labeled 3T3 cells, also color coded by *z* depth. (**B**) *Left*, A similar fluorescence micrograph of assembled pDPAC slides after patterning with a 2×2 flasher design (specifying 3T3 patterning sites using G and MDCK sites using F ssDNAs) and stained with anti-G- and anti-F-FITC probes. *Right*, confocal fluorescence z-projection of a cell-ECM composite kinomorph imaged immediately after release into media. *Insets*, detail of mask design, and corresponding fluorescence *z*-projections of DNA features and H2B-FP-expressing 3T3/MDCK cells.

Having built our first kinomorph design, we studied its behavior in culture. An example flasher kinomorph folded in cell medium over 36 hours into a set of crease networks that had 137 of 142 mountain creases in the expected orientation (96%) compared to the corresponding simulated origami (Fig. S4, Movie S5). Of the properly oriented mountains, 131 (96%) were approximately flat folded (i.e. ECM leaflets on opposite sides of a crease fully contacting each other), creating possible tubule niches that present a basal ECM surface to resident cells in all radial directions within the crease. Indeed, the originally printed 3T3 cells appeared to distribute along creases after an initial phase of traction between clusters transmitted through the gel, in keeping with previous observations in simpler self-folding tissues (19). Based on these observations, we concluded that large self-folding kinomorphs can be successfully programmed to undergo a predictable developmental trajectory that mimics the ECM compaction and leafleting aspects of kidney collecting duct morphogenesis.

With these basic dynamics in place, we sought to add in a kidney epithelial MDCK cell population and study its time-dependent behavior. We therefore created a 2×2 tessellated flasher kinomorph patterned with 3T3s in addition to clusters of MDCKs along gel regions associated with programmed mountain folds (Fig. 3B). Cell populations could be patterned in series with little cross-reactivity using orthogonal ssDNA strand pairs (Fig. S5). In this scheme, we used F/F’ to pattern MDCKs and a second strand pair G/G’ to pattern 3T3s in the 2×2 flasher design. We hypothesized that the engineered crease compaction local to these MDCK clusters would direct them to elongate and fuse as long-range branched tubule networks. 2×2 flashers that were patterned and placed in culture folded radically within the first 20 hr of culture (Fig. 4A, Movie S6) – an example kinomorph had all 142 mountain creases (100%) in the programmed orientation and approximately flat-folded. However, kinomorphs did not progress through even more radically folded shapes predicted by the origami simulator (Movie S7), because even when creases were flat folded, the crease networks appeared to become mechanically frustrated due to their thickness relative to the infinitely thin model sheets. Even so, we concluded that the presence of MDCKs does not interfere substantially with the kinomorph folding program.

**Fig. 4.**
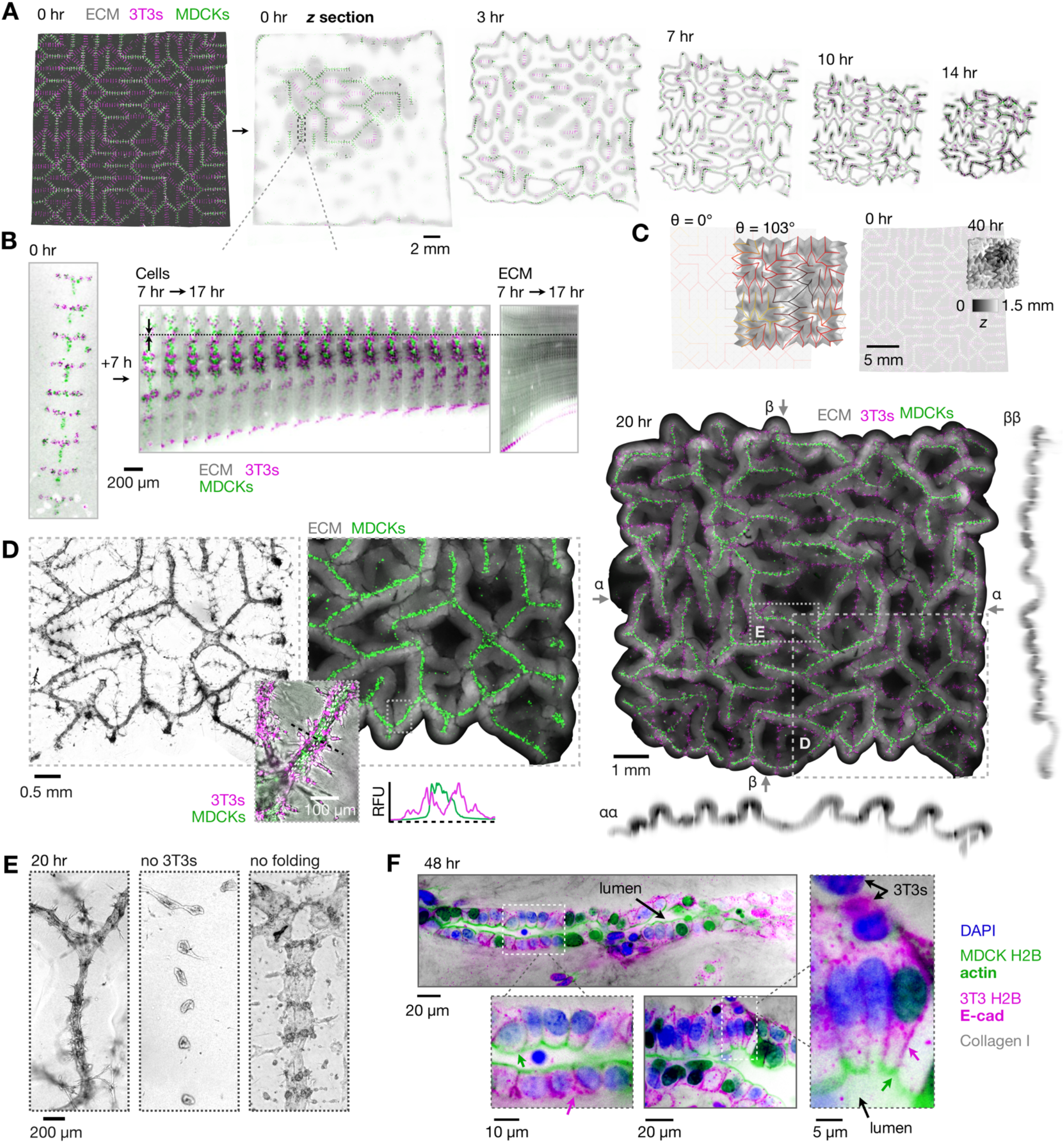
Kinomorphs direct tubule network assembly and structural maturation. (**A**) Confocal fluorescence micrographs from a time-lapse experiment taken at an approximately mid-plane *z* section during kinomorph compaction and folding in culture. (**B**) *Left*, fluorescence micrograph of cells patterned at a single crease. *Middle*, a kymograph of the crease over a period of 7-17 hrs after the kinomorph was placed in culture. Note axial compaction of the cells towards a stationary position in the image (dotted line). *Right*, a kymograph of an ECM region immediately adjacent to cells within the same crease, showing a similar axial compaction of ECM. (**C**) *Left*, output from an origami simulation chosen to match the degree of folding (*SI Text*) of the kinomorph after 40 hr in culture (shown at right as a 3D rendering shaded by *z*-height). (**D**) *Right*, an average z-projection and xz/yz sections of the kinomorph after 20 hr in culture. *Left*, phase and fluorescence micrographs of a region of interest showing MDCK tubules. *Inset*, relative fluorescence traces across a tubule section show basal 3T3 localization. (**E**) Further detail of another region of interest (*left*) compared to non-folding controls due to a lack of 3T3s (*middle*) or physical adhesion to the culture substrate (*right*). (**F**) 40x immunofluorescence micrographs of two regions of interest at creases stained for DAPI (nuclei), F-actin (apical membrane, green arrows), and E-cadherin (E-cad, at lateral cell-cell junctions, magenta arrows) after 48 hr in culture. Note that nuclear fluorescence in the F-actin and E-cad channels overlap with H2B-FP signals.

Given a successful overall folding trajectory, we next used spatial registration of confocal time-lapse images to determine the dynamics of MDCK cell clusters at single creases during the folding process (Fig. 4B, Movie S6). Individual creases significantly compacted in length along their long axes during the first ~20 hr of folding, during which time neighboring MDCK and 3T3 cell clusters fused into continuous tracts that spanned paths through the crease tree of up to ~3 cm. Axial strains of individual creases from the perspective of MDCK cells were −59% ± 9.6% CV after 20 hr, and −51% ± 16% CV (*n* = 5) from the perspective of the surrounding ECM, showing that traction-based compaction of the ECM by cells was the predominant cause of cluster fusion. This compaction along creases led to a much greater than expected change in cross-sectional area of kinomorphs (9.0-fold ± 11% CV, *n* = 3) relative to origami simulator models (1.3-fold) at a similar degree of folding (Fig. 4C, *SI Text*). The simulations therefore appeared to match kinomorph behavior if an extra isometric scaling factor (i.e. uniform shrinkage) was added. This is likely because maintaining edge length is one of the solver goals associated with the origami simulator, whereas no such limitation exists for kinomorphs. Kinomorph compaction led to a 3.8-fold ± 15% CV reduction in the overall volume of ECM (*n* = 3), implying a total collagen I-Matrigel protein concentration upwards of 25 mg/ml on average, and likely higher concentrations local to creases that are the foci of cell mechanical activity. These data together confirmed that MDCK cell fusion into tubules also occurred in engineered creases, similarly to what we had observed for passive crease formation at the edges of floating gels.

Since cell clusters appeared to fuse during kinomorph folding, we next wondered what impact this had on cellcell spacing and self-organization. The result of axial crease compaction and hence global volumetric compaction of ECM is a considerable increase in cell density from the as-patterned state – MDCK cells were initially distributed along prospective creases at an average spacing of ~30 μm, and condensed down to a spacing of 9.0 μm ± 12% CV (*n* = 10) after 40 hr. This spacing is approximately equivalent to the ~8 μm cell spacing seen in E18 mouse collecting ducts (28), and far closer than the ~17 μm spacing that would be expected purely due to cell proliferation over this time. In addition to axial crease compaction, the tight, ordered spacing of cells could be reinforced by axial cell migration along creases, and/or axially-biased cell intercalation (29) – mechanisms that we are currently examining. Besides compact cell spacings, MDCK tubules in creases showed another sign of self-organization – intriguingly, 3T3 fibroblasts appeared to sort concentrically around (i.e. basally to) MDCKs as creases began to adopt flat-folded conformations (Fig. 4D).

We next asked if axial crease compaction was necessary for these self-organization phenotypes. In “no 3T3” control kinomorphs that lacked 3T3s, MDCK cell clusters instead condensed into individual spheroids that either lumenized or protruded into the ECM after 24 hr (Fig. 4E, Fig. S6). In “no folding” control kinomorphs that were adhered to the culture substrate such that ECM deformation at creases was limited, MDCKs and 3T3s spread to form disorganized 2D networks or sheets. We therefore concluded that engineered ECM compaction during flasher folding was necessary for MDCK cluster fusion into continuous 3D tubules along creases that had in vivo-like cell densities and sorting states.

Having characterized the macro- and crease-level behavior of 2×2 flasher kinomorphs, we sought to determine if crease micro-environments were suited to structural maturation of MDCK tubules. We therefore assayed for apico-basal polarization and lumen formation, to determine if tubule cells formed properly localized cell-cell junctions and apical interfaces. Approximately 50% of the length of MDCK+ creases had single-cell columnar MDCK tubules with properly localized F-actin and E-cadherin polarity markers and 3T3s distributed along the basal-ECM interface (Fig. 4F). These tubules were ~30-60 μm in diameter, similar to E12-16 mouse embryonic kidney collecting duct diameters (~50 μm) as well as adult human renal tubules (~40 μm), and collecting ducts (40-100 μm) (30, 31). Remarkably, these tubules formed along engineered kinomorph creases of specified geometry at 10-to-20-fold finer spatial scales than those recently produced through non-dynamic 3D bioprinting approaches (6), and on the scale of the smallest homogeneous collagen filament diameters printed to date (~20 μm) (7). They were also commonly lumenized with a visible lumen cavity or alternatively, with apical cell surfaces pressed against each other (Fig. S7A). Finally, collagen I fibers aligned along the basal surface of these tubules (Fig. S7B), perhaps implying an intimate connection between physical ECM cues at compacted creases and epithelial self-organization there. We concluded that around half of the length of MDCK tubules self-assembled at programmed kinomorph creases showed evidence of structural maturity.

The balance of MDCK+ kinomorph crease areas either had multiple cell layers without clear apicobasal polarity and/or no single apical interface after 48 hr, or were associated with open “atria” typically at tri-fold junctions where adjacent ECM layers were not fully adhered (Fig. S7B). Although we did not culture kinomorphs beyond 72 hr in this first study, we hypothesize that the multiple cell layer tubules could be expected to resolve by a process of lumenization by anoikis, known as cavitation, rather than by the more rapid hollowing associated with cell polarization, apical exocytosis, and membrane separation (13). The atria, on the other hand, could perhaps be sealed off by embedding folded kinomorphs in a second hydrogel layer. Atria also present an intriguing opportunity to interface kinomorphs with other locally self-organizing tissues on the ~200-500 μm length-scale that would lend even finer-scale cellular structure, such as kidney organoids.

## Discussion

Tissue engineers are currently grappling with reconstituting collective cell functions up to the organ-scale. Two promising approaches include 3D printing and organoids, which represent very different conceptual models. The first model is top-down construction, while the second is cellular self-organization guided by reconstituted embryonic cues. Our work attempts to bridge these models with a third option, namely, synthetic reconstitution of developmental motifs in organogenesis.

This new way of building is fully compatible with a range of existing synthetic strategies, including 3D printing. Indeed, researchers are beginning to quantify emergent material dynamics driven by cell behaviors within living bio-inks (32, 33). These dynamics include compaction of relatively loose printed ECMs by resident cells, which can cause shrinkage, fracture, and shape change. Many of these have developmental counterparts including mesenchymal condensation, apical constriction, and cell intercalation (19, 34). Thus, there is an emerging opportunity to understand and then to actively engineer a particular tissue endpoint by manipulating developmental dynamics.

To this end we developed kinomorphs, dynamic tissue scaffolds with which a particular developmental trajectory can be imposed on an initial cell population. We began by creating a high-throughput cell patterning technology that allowed us to extend patterning to >10^7^ spatial sites in total fabrication times of ~2 hr and with single-cell precision. We then mimicked kidney collecting duct morphogenesis by controlling the compaction geometry of ECM sheet into leaflets according to origami design principles. This created networks of crease sites that drove the local self-organization of kidney epithelial tubules.

Our approach here has several limitations. Firstly, we must investigate a more in vivo-like range of cell types and assemblages, for example by integrating primary mouse kidney cell types or human induced pluripotent stem cells at more relevant developmental stages such as ureteric progenitors (11). Our tubule trees, despite spanning cm, are not fully organized along their entire lengths. Vascularization sufficient to guarantee cell viability over long time periods is a challenge for all tissue engineering approaches, although kinomorphs are currently built such that all cells are never >300 μm from media in the mature folding state. Finally, constraining cell proliferation and motility may be a concern for longer culture periods in order to ensure that tissues arrive at a consistent “stopping point”.

Kinomorphs could be extended in a number of ways to achieve more nuanced spatial and compositional structure. Firstly, the ECM scaffold could be engineered for lower thickness, controlled degradation (35), and controlled remodeling by cells (36), to enable tissue structures to pack closer together or to “uncage” new niches over time. Indeed, at 48-72 h timepoints, 3T3 fibroblasts in kinomorph leaflets appeared to extend radially into the surrounding ECM. Such secondary cell behaviors point towards the potential for further finer-scale colonization of the ECM, which could potentially be sculpted using light- or mechanically-actuated biomolecule release peripheral to tubule niches (37).

The kinomorph strategy could also be extended to a 3D analogue, for example, by building networks of filaments that could compact and arrive at a predictable shape, connectivity, and geometry of between-filament interfaces. Such structures could have far higher local cell and ECM concentrations than those achievable in static scaffolds. Kinomorph atria and other geometric niches that emerge could be integrated with organoid technologies to achieve meso-scale (>cm-scale) connectivity. These efforts would build upon the spatial control currently achievable with “assembloid” (38, 39) or cell-material interface (40) strategies (Movie S8). Furthermore, the complementary crease networks available on the other side of kinomorph sheets could be a natural host to engineered vascular beds. More generally, our pDPAC approach should enable multiplexed co-patterning of different cell types to mimic compositional gradients in the kidney. Finally, kinomorphs could host engineered cell populations that activate synthetic cell-cell signaling circuits to control differentiation or spatial cell sorting (41).

In summary, we believe that kinomorphs will serve as a customizable chassis for developmental engineering efforts. Kinomorphs point the way towards an overarching goal to gain engineering control over tissue development at the full range of length-scales necessary to reconstitute entire organs.

## Materials and Methods

### Cell lines and culture

NIH 3T3 mouse embryonic fibroblast cells and Madin-Darby canine kidney epithelial cells were tagged with H2B-fluorescent proteins and cultured at 37°C and 5% CO_2_. See *SI Appendix* for full experimental methods.

### Fabrication of photoactive DNA-programmed assembly of cells (pDPAC) substrates

Glass microscope slides were functionalized with methacrylate groups and used as substrates for polymerization of 30 μm-thick photoactive polyacrylamide gel sheets. See *SI Appendix* for full experimental methods.

### pDPAC photomask design and fabrication

5”x5” chrome-on-quartz mask blanks were fabricated in a cleanroom facility by direct writing to expose a 0.5 μm-thick layer of IP3500 positive photoresist using a DWL 66+ laser lithography system (Heidelberg Instruments), developing in CD-26 solvent, and etching.

### Hardware

Jig parts were designed in Rhino 3D modeling software (Robert McNeel & Associates) and custom CNC-milled in 6061-T651 aluminum (ProtoLabs). Jig halves were bolted together using #6-32 x ¾” machine screws and #6-32 wingnuts (McMaster-Carr).

### ssDNA photolithography on pDPAC substrates

pDPAC polyacrylamide gels were impregnated with ssDNA oligos and sandwiched against photomasks before exposure with 254 nm light to attach DNAs. See *SI Appendix* for full experimental methods.

### Lipid-ssDNA labeling of cells

Cell lines were incubated with lipid-DNA conjugates to passively label them with adhesive DNA strands. See *SI Appendix* for full experimental methods.

### Assembling pDPAC slides and cell patterning

pDPAC slides were spaced apart with a gasket and assembled as a sandwich to form a microfluidic flow cell. Lipid-DNA-labeled cell populations were then introduced and attached to pDPAC slides within the flow cell, and then embedded in a collagen 1-Matrigel ECM hydrogel prior to releasing it into culture. See *SI Appendix* for full experimental methods.

### Fluorescence microscopy, immunofluorescence, and image analysis

pDPAC slides, attached cells, and kinomorphs were analyzed by live fluorescence microscopy and immunofluorescence microscopy using confocal or widefield microscopes. Image analysis was performed in ImageJ/FIJI software (42) and Zerene Stacker software (Zerene Systems). See *SI Appendix* for full experimental methods.

### Origami simulation and similarity analysis

A custom origami simulator was built in Rhino Grasshopper (Robert McNeel & Associates) using Kangaroo2 physics (Daniel Piker). 3D model and kinomorph objects were spatially compared in MeshLab software (43). See *SI Appendix* for full experimental methods.

### Kidney collecting duct and origami tree patterns

We generated the example duct branching pattern for illustrative purposes in Fig. 2 by running a simple Monte Carlo simulation in Matlab (MathWorks) on published tubule tip termination probabilities (25) and manually drawing the resulting tree. The simulation pseudocode is: choose a tip, find tip termination probability *p* at the current generation, generate random number *n* in [0,1], if *n* < *p*, terminate tip, if not, branch tip; repeat until all tips are terminated. Corresponding trees for flasher origamis were drawn by hand from mountain crease networks, ignoring short-circuits where they arose.

### Flasher origamis and translation into pDPAC patterns

The “flasher supreme” origami pattern was originally designed and implemented in paper by Jeremy Shafer (44). 2×2 and 3×3 tessellated designs were inspired by models extended from the Shafer design by YouTube user GreenArt4. Crease block patterns were drawn as binary images in FIJI to denote pixels at which features would be placed, and then read into MATLAB and assembled using a custom script. The resulting matrix of features was then translated into a DXF file using the DXFLib package in MATLAB (Grzegorz Kwiatek).

## Data and Materials Availability

All processed data necessary for interpretation of the study are contained between the article and the *SI Appendix.* Raw data such as fluorescence micrographs are available on request. AutoCAD files containing designs for flasher masks and PDMS gasket, a Rhino file containing 3D plans for jig components, Rhino Grasshopper flasher origami simulations, and MATLAB code for generating mask designs from crease blocks are available on figshare at https://doi.org/10.6084/m9.figshare.9751661.v1

## ACKNOWLEDGMENTS

We thank L.J. Bugaj for H2B-FP constructs and 3T3 cells, and L. Beck and A. Raj for MDCK cells. We appreciate K. Susztak and M. Little for discussions on kidney biology and organoids. We thank D. Patterson and Z. Gartner for test aliquots of lipid-DNAs and assistance with ordering full custom syntheses. This work was carried out in part at the Singh Center for Nanotechnology, which is supported by the NSF National Nanotechnology Coordinated Infrastructure Program under grant NNCI-1542153. This work was partially funded through an NIH MIRA grant (R35GM133380) to A.J.H.

