## Supplemental Text for "Kinomorphs: Shape-shifting tissues for developmental engineering"

### SI Text

**Definition of degree of folding.** We sought to define a comparative method to qualitatively match different folding states between model origamis and kinomorphs. We suggest that models/kinomorphs at similar "degrees of folding" would have approximately matched angles $\phi$ between adjacent creases in *z* projections of their 3D shapes (Fig. S8). Thus, we typically visually matched a kinomorph to a family of model origamis generated at different target rest angles θ and chose the closest match that minimized $\sum_i(\phi_{i,model}-\phi_{i,exp})$.

### SI Materials and Methods

**Cell lines, culture, and fluorescent labeling.** NIH 3T3 mouse embryonic fibroblast cells (male, gift of Lukasz Bugaj, UPenn) expressing histone H2B fused to venus fluorescent protein via pHR-pGK-RBD-H2B-VFP (gift of Lukasz Bugaj, UPenn) and Madin-Darby canine kidney epithelial cells (MDCK-II, female, gift of Arjun Raj, UPenn) expressing H2B-iRFP via pLentiPGK-DEST-H2B-iRFP670 (gift of Lukasz Bugaj) were cultured on polystyrene plates and flasks (Corning) at 37°C and 5% $CO_2$. 3T3s were maintained in Dulbecco's modified Eagle's medium (DMEM, ThermoFisher Scientific 11965084) with 10% bovine calf serum (GE Healthcare Life Sciences SH30087.03), and MDCKs were maintained in minimum essential medium (MEM, with Earle's Salts and L-glutamine, Corning 10-010-CM) and 10% fetal bovine serum (Corning 35-010-CV). Where indicated, the parental MDCK cell line was fluorescently labeled with membrane permeable dyes CellTracker Green or Deep Red according to manufacturer protocols (ThermoFisher Scientific C7025, C34565).

**Fabrication of photoactive DNA-programmed assembly of cells (pDPAC) substrates.** 30 μm-thick polyacrylamide gels were fabricated on methacrylate-functionalized glass slides against SU-8 shims anchored to hydrophobic silicon wafers using standard photolithography methods roughly according to previously published methods (1). SU-8 2025 photoresist (Y111069, MicroChem) was spun (Instras Scientific, SCK-300P) onto mechanical-grade silicon wafers (University Wafer) to a thickness of 30 μm according to manufacturer guidelines and exposed to 365 nm UV light (ThorLabs M365LP1, ACL7560U, SM3V10) at 10 mW/cm$^2$ (Thorlabs PM100D light meter with S120VC 200-1100 nm probe) for 30s under a Mylar mask printed at 20,000 d.p.i. (CAD/Art Services). Exposed wafers were then processed with SU-8 developer solution (Y020100, MicroChem), and alternating washes of isopropanol and acetone. The mask was designed such that SU-8 shims ran along the long edges of the glass slide, setting a 30 μm gap between the slide and the bare silicon. Wafers were silanized by vapor-deposition of 2 ml of the hydrophobic silane dichlorodimethylsilane (DCDMS, 440272, Sigma-Aldrich) for 1 h *in vacuo*, washed thoroughly with deionized (DI) water, and dried under a stream of compressed air immediately before use. Silanized wafers were robust to reuse after gel fabrication after rinsing wafers with 0.1% Triton in DI water and DI water in excess of 20 times.

Plain pre-cleaned 75 x 50 mm glass microscope slides (Corning 2947-75X50) were rinsed in 0.1% Triton to remove surface grease, and either dried (for the bottom slide) or laser etched to create through-holes (for the top slide) using 20-30 passes of a 50 W etching laser at 100% power, 15% speed, 350 pulses per inch (VLS3.5, Universal Laser Systems). Slides were then silanized to establish a self-assembled surface monolayer of methacrylate functional groups according to standard protocols (Hughes 2010). Silanized slides were placed face-down on wafers and manually aligned to the SU-8 shims. Gel precursor solutions were 4%T (w/v total acrylamides), 2.7%C (w/w of the cross-linker *N*,*N*-methylenebisacrylamide) from a 30%T, 2.7%C stock (A3699, Sigma-Aldrich); 3 mM benzophenone-methacrylamide (N-[3-[(4-benzoylphenyl) formamido]propyl] methacrylamide, BPMAC) from a 100 mM stock in DMSO, 0.06% SDS (161-0301, Bio-Rad), 0.06% Triton X-100 (BP151, Fisher), 0.05% ammonium persulfate (APS, A3678, Sigma-Aldrich), and 0.05% tetramethylethylenediamine (TEMED, T9281, Sigma-Aldrich) in 25 mM Tris, 192 mM glycine, pH 8.3, (1x from a 10x stock, 161-0734, Bio-Rad). BPMAC can be synthesized relatively simply as previously described (2), however, we used BPMAC from a commercially synthesized lot (PharmAgra). A partial precursor mixture containing acrylamides and buffer was degassed (15-337-411, Fisher) for 1 min *in vacuo* immediately before the addition of detergents (SDS, Triton), BPMAC, and finally, polymerization initiators (APS, TEMED). The full precursor was then injected into the gap between the glass slide and silicon wafer using a standard 200-µl pipette. After allowing ~30 s for precursor to wick through the gap, polymerization was allowed to continue for an additional 20 min. Gel-fabricated glass slides were wetted at their edges using 2 ml of TAE buffer (40 mM Tris, 20 mM acetic acid, 1 mM EDTA, pH 8.3) and carefully levered from wafers using a razor blade. Fabricated slides could be rinsed in TAE, dried under a stream of compressed air and stored at 4 °C for at least 2 weeks before use without loss of photocapture properties.

**ssDNA photolithography on pDPAC substrates.** pDPAC slides were patterned with 5'-polyT$_{20}$-X$_{20}$-3' ssDNA oligos (IDT) against custom chrome-on-quartz masks in a glove box (Bel-Art, H50028-2001) containing a medical-grade nitrogen atmosphere. X$_{20}$ for strand "F" is 5'-AGAAGAAGAACGAAGAAGAA-3', for strand "G" is 5'-AGCCAGAGAGAGAGAGAGAG-3', and for strand "A" is 5'-ACTGACTGACTGACTGACTG-3'. Dried pDPAC gels were rehydrated in the glove box with 150 µl of degassed oligo at 2.5 mM in TAE by drawing a bead across the slide using the edge of a 50 x 64 mm glass coverslip (Electron Microscopy Sciences, 71881-90). After evenly distributing the liquid bead, another 150 µl of oligo solution was added to the mask, and the pDPAC slide was laid face-down on the mask starting from one of the long edges, laying the slide down carefully to avoid introducing bubbles. Excess fluid was withdrawn using a kimwipe. The slide-mask sandwich was then removed from the glove box and either immediately exposed to 254 nm UV light (Stratagene) at ~7 mW/cm$^2$ for 2 min, or if alignment to a pre-existing oligo pattern was necessary, the sandwich was inspected and manually aligned using an upright wide-field fluorescence microscope (Nikon Ts2-FL with LED-based epifluorescence) prior to exposure (Fig. S9). Registration marks needed for alignment of additional ssDNA strand(s) were stained using 1 µM of the appropriate anti-X$_{20}$-FITC probe (IDT) in TAE for 20 min and washed in a 15 cm petri dish with 15 ml TAE twice for 10 min per wash. After 254 nm exposure, the slide was carefully levered from the mask using a razor blade after wetting its edges using 2 ml of 0.1% SDS in TAE buffer. The slide was then transferred to 15 ml of 0.1% SDS in TAE in a 15 cm petri dish and washed for 10 min, followed by two similar washes in TAE. The slide was then dried under a stream of compressed air if another oligo was to be patterned, or alternatively, briefly washed in DPBS ($Ca^{2+}$/$Mg^{2+}$-free, Corning 21-031-CM) and dried prior to cell patterning.

**Lipid-ssDNA labeling of cells.** Cell lines were lifted from T150 culture flasks by incubating with 0.05% (3T3s) or 0.25% Trypsin (MDCKs) (ThermoFisher Scientific, 25300054 and -56) for approx. 10 min at 37°C before resuspending in cell media, centrifuged (200 g, 3 min, 4°C), and washed twice by resuspending in 10 ml DPBS and pelleting cells. Washed pellets were resuspended in 1.5 ml eppendorf tubes to 100 µl DPBS and labeled with lipid-DNAs (custom syntheses, OligoFactory, Holliston, MA) by first adding “universal anchor” DNA 5’-TGGAATTCTCGGGTGCCAAGGGTAACGATCCAGCTGTCACT-C24 lipid (lignoceric acid)-3’ to 1.5 µM final concentration from a 100 µM stock in DI water, followed by 1.5 µM final of “universal co-anchor” DNA 5’-C16 lipid (Palmitic acid)-AGTGACAGCTGGATCGTTAC-3’, followed by 1.5 µM final of “adhesion strand” DNA 5’-CCTTGGCACCCGAGAATTCCA-poly$T_{20}$-$Y_{20}$, where $Y_{20}$ is the reverse complement of the $X_{20}$ sequence patterned on the pDPAC slide to which cells were to be adhered (3). Each oligo was added in succession to the 100 µl reaction, with 5 min incubation steps between each addition under gentle agitation on a vortex set at very low speed (approx. 5 Hz). After adding the series of 3 oligos, cells were washed in 3 x 1 ml volumes of DPBS to remove free DNAs. We found that cell capture efficiency was improved by labeling no more than approx. 5 x $10^6$ cells (approx. 1 x T150’s worth) per labeling reaction. If more cells were to be labeled, we split them into an appropriate number of parallel reactions. At the end of labeling and washing, cells were pooled and resuspended to 600 µl prior to introducing to the pDPAC slide flow cell.

**Assembling pDPAC slides and cell patterning.** If not already done, registration marks on the top and bottom pDPAC slides were labeled with anti-$X_{20}$-FITC probe as described. The bottom slide was placed onto a flat surface and a craft-cut (Silhouette Cameo 3), 0.01”-thick PDMS gasket (SSP Inc., M823), was centered on the slide and pressed down gently with forceps. After transferring the slide and gasket to the aluminum jig base, 3-5 ml of DPBS was pooled on top of the gel layer within the gasket, and the top pDPAC slide laid down from one short edge to the other in order to avoid trapping bubbles in the space. The top of the aluminum jig with a ⅜” silicone rubber gasket (McMaster-Carr, 1460N27) was then placed and lightly tightened against the slide sandwich before the top and bottom slides were manually aligned on the inverted fluorescence microscope. The rubber gasket here was cut to shape with a razor blade to match the size of the opening in the top part of the jig. Jig wingnuts were then progressively tightened, working in a circular pattern until a light finger-tight tension was attained evenly across the jig.

With the flow cell jig placed on ice, 500 µl of cell suspension was introduced to flow cells by pipetting on top of one of the sets of through-holes and removing and discarding fluid from the opposite set of through-holes using a p200 pipet. We found that approx. 15 x $10^6$ cells (3 x T150’s worth) were needed to form a confluent blanket within 2” x 3” slide flow cells during cell patterning. The remaining 100 µl of cell suspension was cycled three times from outlet wells to inlet wells every 2 min, during which time cell settling and adhesion to DNA spots was achieved. A similar process set of cycles and pauses was performed to achieve cell patterning to the top pDPAC slide, except the jig was flipped upside-down during each 2 min pause so that cells could contact it. Cells were washed out of the flow cell using around 2-3 ml DPBS placed at the inlet wells and removed from the outlet wells. A further round of cellular assembly was used to generate clusters of 5-8 cells at each DNA feature in the case of 3T3s. If lipid-ssDNAs on cell membranes and photo-patterned ssDNAs were to be imaged *in situ*, we introduced 10x SYBR Gold in DPBS (from a 10,000x stock, ThermoFisher Scientific, S11494) for 5 min and washed with DPBS prior to imaging. Otherwise, 1 ml of liquid ECM gel precursor was introduced and the jig placed at 37°C for 40 min to set the precursor. The ECM gel here consisted of a neutralized composite of fluorescently-labeled collagen I fibers in Matrigel. To prepare this precursor, we made "Solution A": 1 µl 10M NaOH and 199 µl 10X DPBS (ThermoFisher Scientific 14200-075). Then "Solution B": 10 µl 1 mg/ml Alexa 555-NHS in DMSO (ThermoFisher Scientific A20009), 350 µl 3.9 mg/ml rat tail collagen I (Corning 354236), 40 µl Solution A. Then after incubating Solution B for 10 min on ice, the final mix was: 280 µl Solution B, 690 µl 8.5 mg/ml growth factor-reduced Matrigel (Corning 354230), 30 µl TurboDNAse (ThermoFisher Scientific AM2238).

After setting the ECM gel in the flow cell, the jig was gently disassembled and the slide sandwich submerged in cell media at room temperature within a 15 cm x 2.5 cm “deep style” petri dish (Corning 430597) pre-fitted with a base layer of 10 mm-thick 10:1 polydimethylsiloxane (PDMS) silicone rubber (Sylgard 184, Fisher, 50-366-794) containing a 55 x 70 mm recess. A razor blade was used to gently pry apart the glass slides. Kinomorphs bearing patterned cells typically floated spontaneously into the media or could be gently detached from pDPAC slides (often catching at the wells) with a micro-spatula (Fine Science Tools 110089-11). Kinomorphs were then manually dissected using a razor blade. Finished tissues were transferred to a custom incubation chamber previously submerged in the PDMS recess consisting of a 50 x 64 mm glass coverslip (Electron Microscopy Sciences, 71881-90) bonded to a 9 mm-thick piece of PDMS with a 35 x 35 mm cut-out large enough to fit the kinomorph. The coverslip bottom of the chamber was also pre-coated with a thin layer of 1% agarose (Fisher BP1356-500) in PBS, serving as a non-adhesive underlay. For non-folding controls, we omitted the agarose underlay, and large kinomorphs adhered spontaneously during the early stages of incubation, preventing them from folding. Custom incubation chambers were covered with custom 55 x 70 mm lids constructed from laser-cut 1/16”-thick acrylic sheet (McMaster-Carr 8560K171) glued with acrylic cement (Weld-On 4).

pDPAC slides could be recovered after releasing the kinomorph and stained with 10x SYBR Gold in TAE for detailed fluorescence imaging or visual validation of correct ssDNA patterning using a blue light LED transilluminator (IMG-04-03, IO Rodeo Inc.). Jig and gasket components were rinsed in 70% ethanol and reused.

**Quantifying the effect of nucleotide composition on ssDNA patterning.** For the systematic variation of ssDNA nucleotide composition to determine effects on benzophenone-mediated capture to pDPAC substrates (Fig. S1), we used 5’-$X_{24}$-$Y_{21}$-3’ ssDNA features, where $X_{24}$ is a random 24-mer variable sequence composed of different proportions of bases, and $Y_{21}$ (5’-TATCAGGAAACAGCTATGACG-3’) is a constant 21-mer probe binding sequence. This design allowed us to use a single probe sequence (5'-FITC-CGTCATAGCTGTTTCCTGATA-3’) to detect bound DNAs rather than SYBR Gold, which could potentially label different sequences with different affinities and thereby complicate analysis. The 5’-$X_{24}$-$Y_{21}$-3’ sequences were as follows:

A: AAAAAAAAAAAAAAAAAAAAAAAA TATCAGGAAACAGCTATGACG
T: TTTTTTTTTTTTTTTTTTTTTTTT TATCAGGAAACAGCTATGACG
C: CCCCCCCCCCCCCCCCCCCCCCCC TATCAGGAAACAGCTATGACG
½ A, ½ T: TAATTATATATATTATTATAAATA TATCAGGAAACAGCTATGACG
½ C, ½ T: CTTCTCTCTTCTTCTTTTCCCCCC TATCAGGAAACAGCTATGACG
½ A, ½ C: AACCAAACCACCCCACAAAACCCA TATCAGGAAACAGCTATGACG
⅓ A, ⅓ C, ⅓ T: CCAAACTCTATACTATTCACTCTA TATCAGGAAACAGCTATGACG
¾ A, ¼ C: ACAAAACAAAACAAAACAACACAA TATCAGGAAACAGCTATGACG
¼ A, ¾ C: CAACCCAACCCACCCCCCCACCCC TATCAGGAAACAGCTATGACG

**Fluorescence microscopy.** 3D stacks and time-lapses were recorded in an environmental chamber (OkoLab) containing a 37°C and 5% $CO_2$ atmosphere through 4x-20x Nikon CFI60 Plan Apochromat air objectives via tiled confocal microscopy. Images were acquired through Nikon Elements software controlling a Nikon Ti2-E with a Yokogawa CSU-W1 spinning disk and 4-line laser source (405nm, 100mW; 488nm, 100mW; 561nm, 100mW; 640nm, 75mW with fiber output), 2048x2048 Photometrics Prime BSI sCMOS camera, piezo *z* stage, motorized *xy* stage and light distribution. Kinomorph folding time-lapses were typically captured at a single *z* plane using tiled 4x exposures every 20 min.

**Immunofluorescence.** Kinomorphs were fixed in 2% paraformaldehyde for 45 min at room temperature. All pipetting was done carefully to avoid mechanical disruption with the pipet tip. Kinomorphs were washed three times with 100 mM glycine in PBS for 20 min per wash and permeabilized in 0.5% Triton X-100 in PBS for 15 min, all at room temperature; and blocked overnight at 4°C in 0.1% bovine serum albumin, 0.2% Triton X-100, 0.04% Tween-20, 10% goat serum (ThermoFisher Scientific 16210064) in PBS. Kinomorphs were labeled with rabbit anti-E-Cadherin (Cell Signaling Technology 3195) diluted 1:200 in blocking buffer overnight at 4°C and washed three times for 1 hour per wash in blocking buffer. This process was repeated for 1:200-diluted AlexaFluor 647-labeled donkey anti-rabbit secondary antibody (ThermoFisher Scientific A32795), followed by a 15 min incubation in PBS containing 1 µg/ml DAPI (for nuclei) and 0.17 µM AlexaFluor 488-labeled phalloidin (for F-actin, ThermoFisher Scientific A12379), and a final PBS wash. Tissues were then imaged at 20x air or 40x water in coverslip-bottom chambers in FocusClear (CedarLane FC-101).

**Image analysis**. Fluorescence images were manipulated using linear processes such as brightness/contrast, binning, inversion, and *z*-projections (either maximum or average) in ImageJ/FIJI software (4). Kymographs were also generated in FIJI using a montage of image strips in a time-lapse stack. Focus stacking was used to create images that spanned a series of focal points in brightfield/phase image *z* stacks of kinomorphs using Zerene Stacker software (Zerene Systems).

**Origami simulation and similarity analysis.** “2.5D” origami sheet folding was simulated using Kangaroo2 (Daniel Piker), a position-based dynamics solver within the Rhino Grasshopper (Robert McNeel & Associates) algorithmic modeling environment (5). LunchBox (Nathan Miller) and Shortest Walk Gh (Giulio Piacentino) plug-ins are also required. We took a form-finding FEM approach to enable interactive, rapid prototyping, at the expense of high-resolution simulation of matrix deformations. Square unit cells were constructed from quad mesh faces with two diagonals per quad. Unit cells were assembled to model origamis at a scale of 1.4 mm per unit model edge length, balancing spatial resolution and simulation time. All edges were modeled as linear elastic elements with stiffness k and rest lengths equal to their initial lengths (*EdgeLengths* goal function). The edges in the xy plane coincident with valley or mountain creases were specified as active hinges with rest angle θ manipulated by the user (*Hinge* goal function). The *NoFoldThrough* goal function was used to discourage hinge faces from passing through each other. Crease distances from the network “trunk” were computed using the *ShortestWalk* function.

Models constructed in this way had two parameters – the stiffness of edges k, and the active edge rest angle θ. Note that we did not attempt to capture axial crease compaction in the model. k = 0.4 $Nm^{-1}$ was chosen to give models some incidental pliability but to resist excessive edge deformation, such that objects appeared to behave roughly as paper origami models. Origamis were typically simulated for at least 30 values of θ in the range of 0-2π to produce a family of objects at intermediate folding states for each design. Since θ is not directly analogous to tissue folding time in vitro, models intended for comparison to kinomorphs at a given time-point were chosen from object families based on qualitative similarity in degree of folding (*SI Text*). Simulated tissue models were exported from Rhino as stereolithography (.stl) files for Hausdorff distance comparison.

Full 3D finite element models simulating ECM compaction and curvature at the level of cell clusters and simple crease networks up to 31 folds were generated using cuboidal unit cells in Grasshopper as previously described (6) and used here for Hausdorff distance comparison. Hausdorff distances (7) were computed between model meshes and meshes reconstructed from confocal fluorescence microscopy data (see (6)) using MeshLab software (8).

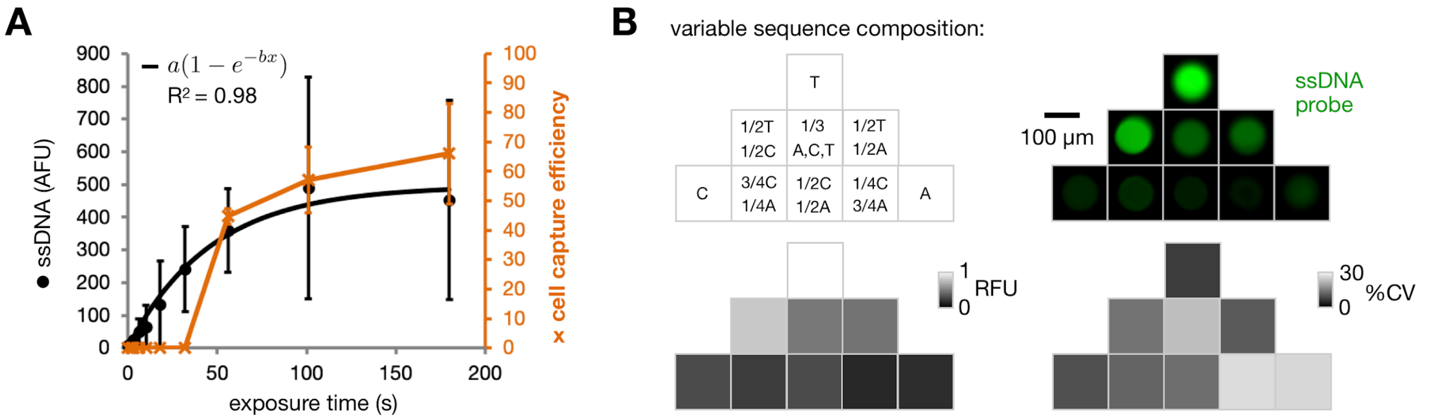


**Fig. S1. ssDNA patterning on pDPAC substrates depends quantitatively on UV dose and nucleotide composition.** (A) *Left axis*, Plot of the amount of 2.5 mM F ssDNA patterned onto pDPAC gels as a function of exposure time to a 254 nm light source (*SI Materials and Methods*), as determined by arbitrary fluorescence intensity (AFU) of SYBR Gold-labeled features imaged by confocal microscopy (± SD, *n* = 3 experiments from an average of 10 features per experiment condition). *Right axis*, capture efficiency of F' lipid-DNA-labeled MDCK cells to features, measured as the % surface area of features covered by attached cells imaged by phase microscopy (± SD, *n* = 3 experiments from an average of 5 features per experiment condition). Note that capture efficiency is limited to a maximum of 91%, which is set by the geometry of hexagonal packing of circles in a 2D plane. (B) *Top*, Example fluorescence micrographs of anti-$Y_{21}$-FITC probe-labeled 5'-$X_{24}$-$Y_{21}$-3' ssDNA features, where $X_{24}$ is a random 24-mer variable sequence composed of the indicated proportions of bases, and $Y_{21}$ is a constant 21-mer probe binding sequence (*SI Materials and Methods*). *Bottom*, heat maps of relative fluorescence (RFU) and %CV across *n* = 3 experiments from an average of 10 features per experiment condition.

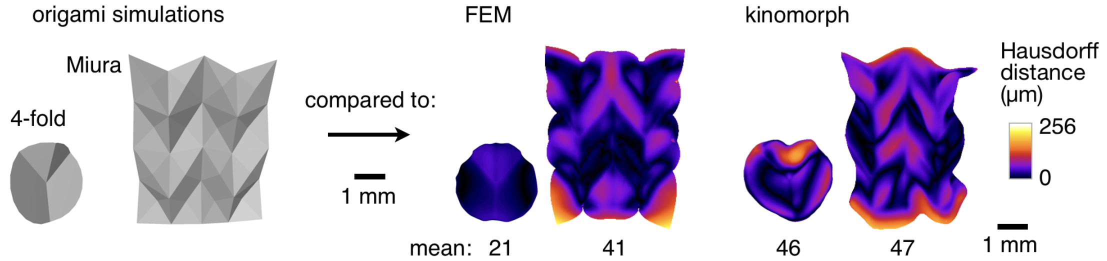


**Fig. S2. Kinematic origami simulations faithfully approximate finite element and kinomorph geometries.** 3D mesh objects for finite element models (FEM) and experimentally produced kinomorphs for 4-fold and Miura origami designs shaded by local Haussdorff distance from origami simulation meshes at similar degrees of folding (*SI Text*). FEM and kinomorph meshes were adapted from (6).

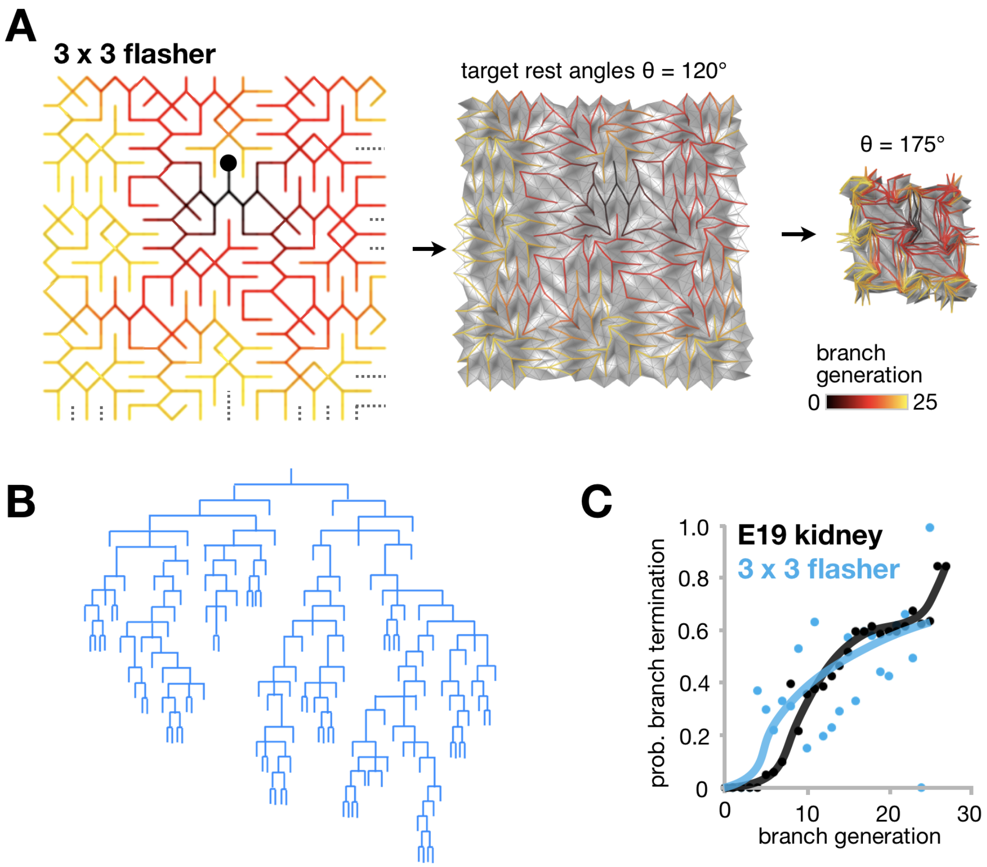


**Fig. S3. Flasher origamis can be tessellated to at least 3x3 without loss of kidney-mimetic branching properties.** (**A**) Origami simulation of 3x3 flasher edge network, color coded by the number of branch generations from a given edge to the indicated center point (filled black circle). (**B**) Branch pattern describing the edge network, ignoring the occasional short circuit. (**C**) Plot of the probability that a flasher edge terminates at a given branch generation from the center point, compared to similar data for embryonic day 19 mouse kidney (modified from (9)).

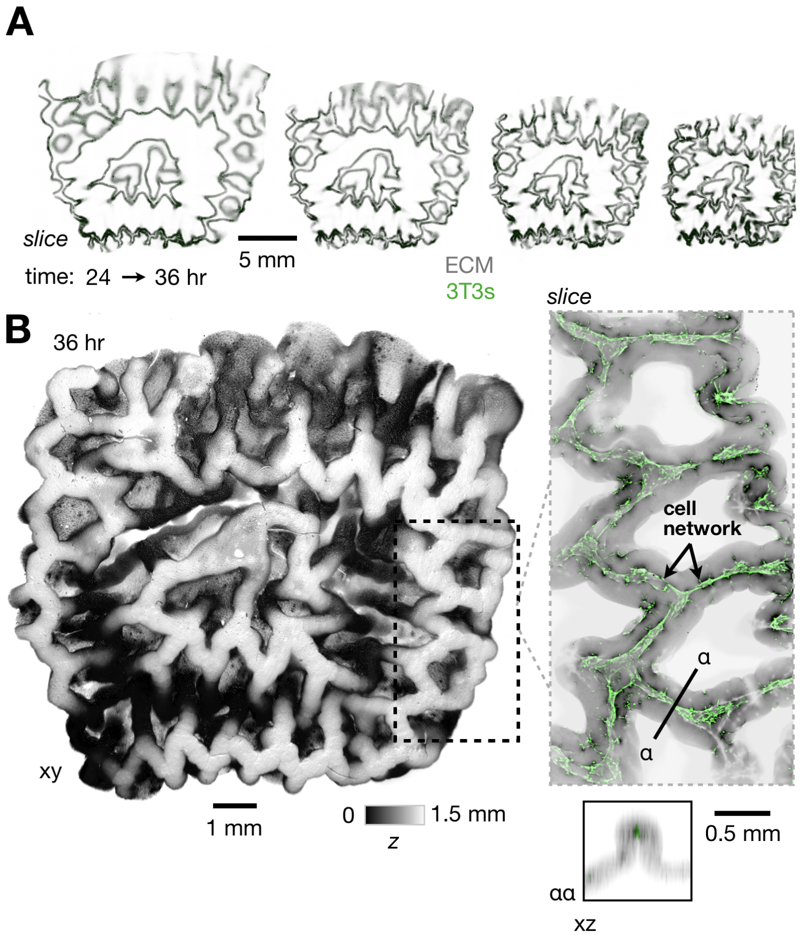


**Fig. S4. Flasher kinomorph demonstrates large-scale engineering of ECM deformation.** (**A**) Confocal fluorescence micrograph frames from a live imaging time-lapse taken approximately at the mid-plane in *z* of a flasher kinomorph built according to the design presented in Fig. 3A, compacting and folding in culture. (**B**) 3D rendering of a confocal fluorescence micrograph stack taken after 36 hr of culture, shaded by *z*-height. *Inset*, z-slice and xz section of the stack at approximately the plane of ridge creases in the kinomorph, showing 3T3 fibroblasts spread along approximately flat-folded creases.

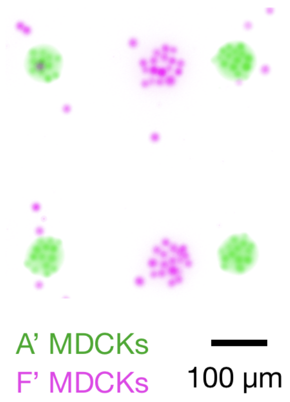


**Fig. S5. Orthogonal ssDNA strand pairs enable independent patterning of cell populations.** Fluorescence micrograph of MDCK cell populations labeled with CellTracker Green (green) or Deep Red (magenta) attached to pDPAC-generated ssDNA patterns via A/A’ and F/F’ ssDNA strand pairs respectively.

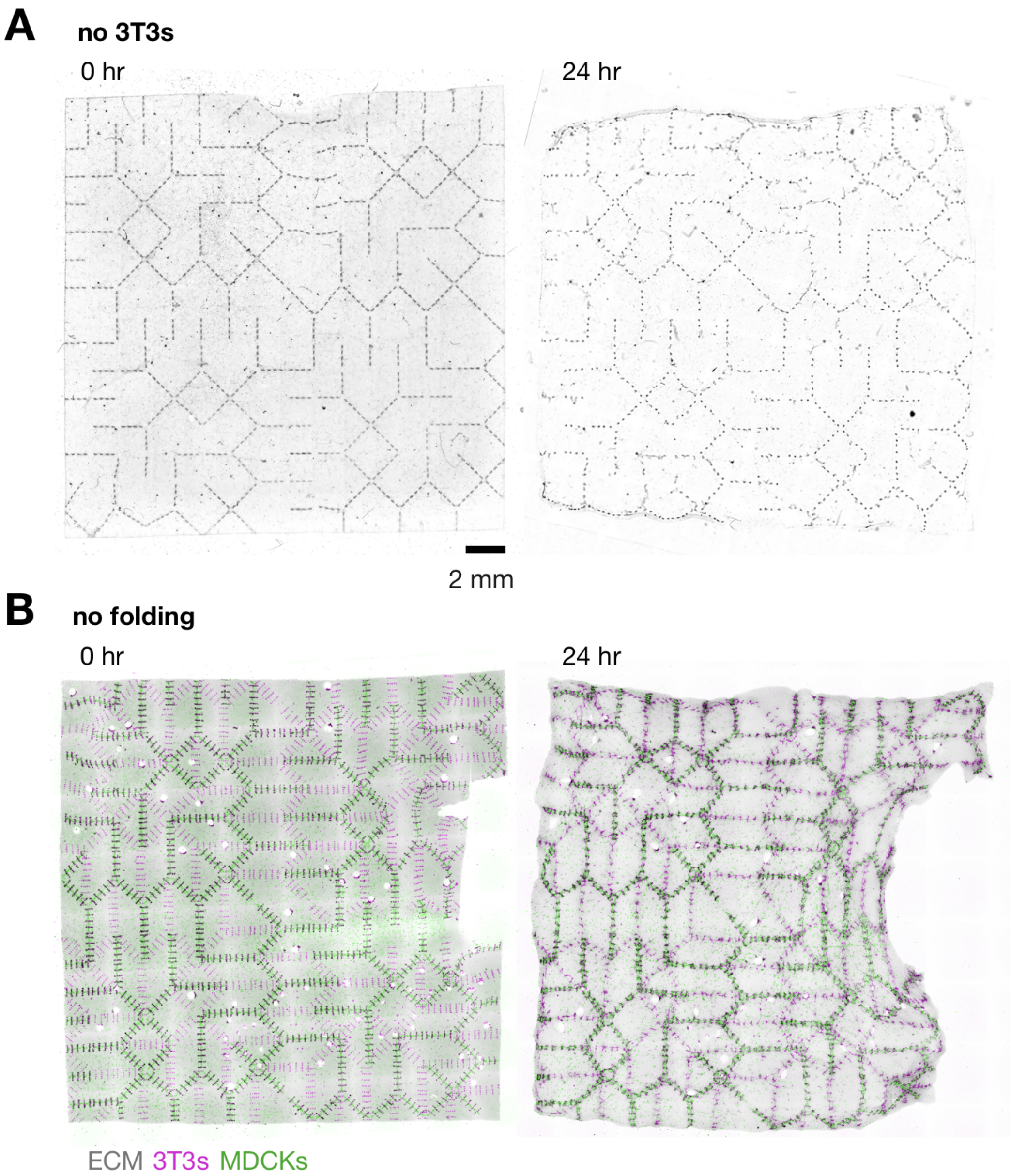


**Fig. S6. Engineered ECM compaction during folding is necessary for 3D tubule formation.** (**A**) Phase contrast micrographs of a 2x2 flasher kinomorph lacking 3T3s immediately after transfer to culture and after 24 hr in culture. MDCK clusters locally assembled into spheroids. (**B**) Confocal fluorescence micrographs for a similar experiment in which both MDCKs and 3T3s were patterned, but where the kinomorph was allowed to adhere to the culture substrate, thereby preventing folding. MDCKs and 3T3s merged and spread as disorganized 2D sheets.

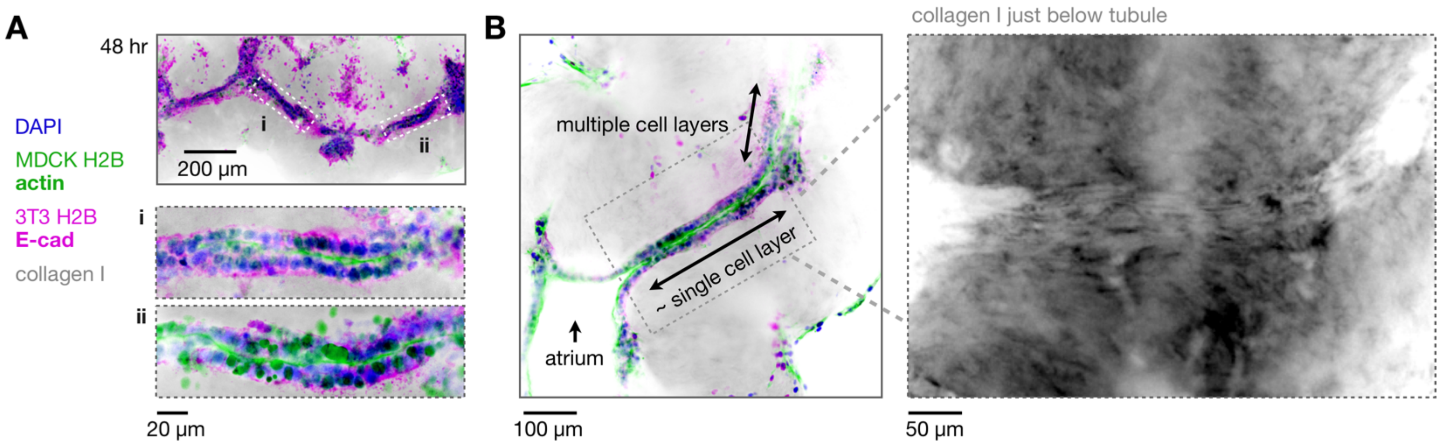


**Fig. S7. Other kinomorph tubule phenotypes.** (**A**) Confocal immunofluorescence micrographs of example 2x2 flasher kinomorph edges at which tubules polarized in single-cell layers, but where apical cell membranes were pressed together rather than fully lumenized. (**B**) *Left*, Confocal immunofluorescence micrograph of a tubule bordered by an open "atrium" at a tri-fold junction where adjacent ECM layers were not fully adhered, and an area with multiple cell layers. *Inset*, Collagen I fiber fluorescence immediately below the tubule, showing fiber alignment along the basal epithelial interface.

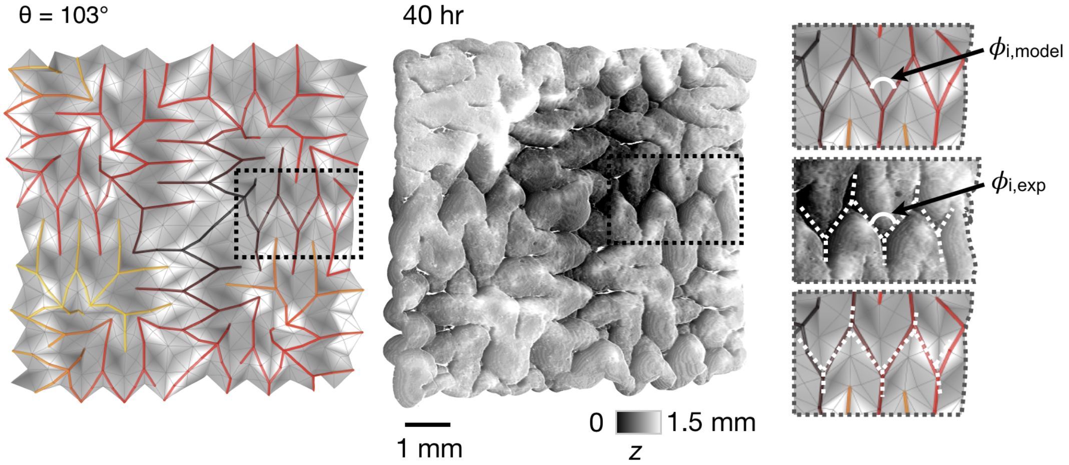


**Fig. S8. Semi-quantitatively matching the degree of folding between model and experiment kinomorphs**. *Left*, 2x2 flasher origami simulation kinomorph with ridge edge network generated at the indicated target rest angle θ. *Right*, corresponding kinomorph imaged after 40 hr in culture, shown as a 3D-rendered confocal fluorescence stack shaded by *z*-height. *Inset*, tracing ridges in the model and kinomorph *z*-projections to demonstrate an adjacent crease angle $\phi_i$. Model origamis were chosen from families generated at different θ values and visually matched to a given experiment kinomorph to approximately minimize $\sum_i(\phi_{i,model}-\phi_{i,exp})$.

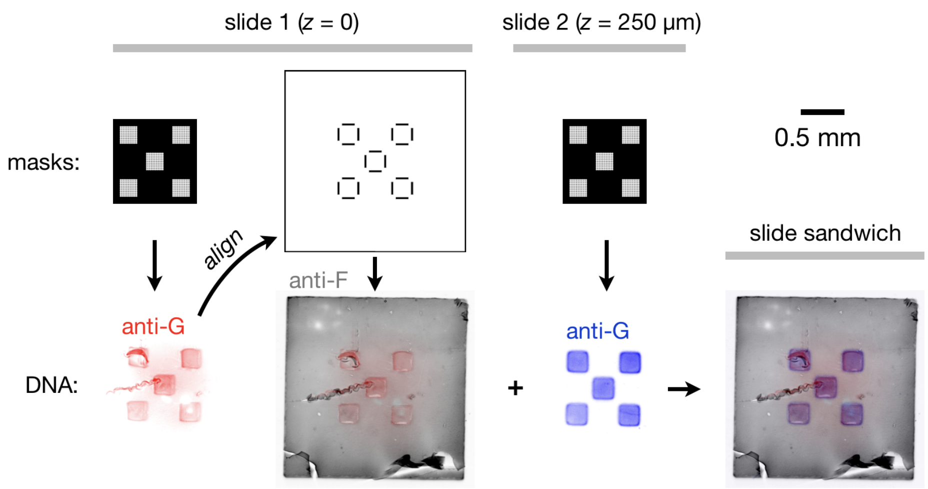


**Fig. S9. Registration marks enable visual alignment of ssDNA patterns on the same pDPAC slide, and on sandwiched slides.** *Left*, mask design, confocal fluorescence micrographs of the resulting G ssDNA pattern imaged using an anti-G-FITC probe, alignment to a second mask design applied to the same pDPAC slide, and resulting 2-plex ssDNA pattern imaged using a second anti-F probe. ssDNA associated with the X-shaped set of squares on the first mask is visually matched to a “window” pattern on the second mask to achieve spatial registration. *Right*, mask design, resulting G ssDNA pattern by confocal, and appearance of overlayed features on the first and second slides assembled into a sandwich and imaged by wide-field fluorescence microscopy.

**Movie S1 (separate file).** Demonstration of compaction and leafleting analogies between (**A**) folding, and (**B**) branching. Folding a sheet with conserved volume compresses individual leaflets against each other, similar to epithelial tubules undergoing branching morphogenesis into an ECM volume that does not grow at the same rate as the epithelial volume.

**Movie S2 (separate file).** *Left*, Confocal fluorescence time-lapse showing collagen (coll.) I dynamics and fusion of CellTracker Green-labeled MDCK cell clusters built by photolithographic DNA-programmed assembly of cells at pliable ECM edges over 5 hr. *Right*, fluorescence micrograph showing tubule formation due to cell cluster fusion after a further 24 hr.

**Movie S3 (separate file).** Animated projections of a confocal fluorescence micrograph z-stack showing SYBR Gold-labeled F ssDNA features on two pDPAC slides sandwiched together at a distance of 250 μm apart in *z*.

**Movie S4 (separate file).** Animation showing flow cell assembly from two 2”x3” pDPAC slides, schematic of ssDNA features on both slides including registration marks used for slide alignment, and a photograph of SYBR Gold-labeled ssDNA features within the slide sandwich held over a blue LED light source.

**Movie S5 (separate file).** (**A**) Confocal fluorescence time-lapse taken approximately at the mid-plane in *z* of a flasher kinomorph built according to the design presented in Fig. 3A, compacting and folding during 24-36 hr after transferring to culture. (**B**) Successive confocal fluorescence z-slices at 50 μm increments over a *z* range of 1.5 mm for the kinomorph at 36 hr of culture. (**C**) 3D rendering of a confocal fluorescence micrograph stack taken at 36 hr, shaded by *z*-height. *Inset*, z-slices at 25 μm increments over a *z* range of 0.6 mm for a region of interest through ridge creases in the kinomorph, showing 3T3 fibroblasts spread along approximately flat-folded creases.

**Movie S6 (separate file).** (**A**) Confocal fluorescence time-lapse taken approximately at the mid-plane in *z* of a 2x2 flasher kinomorph built according to the design presented in Fig. 3B, compacting and folding during 0-17 hr after transferring to culture. (**B**) Successive confocal fluorescence z-slices at 60 μm increments over a *z* range of 2 mm for the kinomorph at 20 hr of culture. (**C**) 3D rendering of a confocal fluorescence stack taken at 20 hr, shaded by *z*-height. *Inset*, two regions of interest from the time-lapse in (**A**), showing kinomorph crease compaction and cell fusion.

**Movie S7 (separate file).** Time-lapse of 2x2 flasher origami simulation kinomorph with ridge edge network generated over a range in target rest angle θ = 0-166°.

**Movie S8 (separate file).** A vision for organoid scaffolding using kinomorphs in a series of niches interconnected by tree-like tubule and vascular networks.
